# Mechanosensory cephalic bristles mediate rapid flight initiation in endothermic hawkmoths

**DOI:** 10.1101/2024.02.28.582474

**Authors:** Maitri Manjunath, Chinmayee L. Mukunda, Sanjay P. Sane

## Abstract

Endothermic insects including bees, butterflies, and moths need to warm up their flight muscles before taking flight. For instance, diurnal butterflies bask in the sun to heat their flight muscles, whereas nocturnal hawkmoths display a pre-flight shivering behavior in which small-amplitude wing movements cause flight muscles to warm up, eventually generating large-amplitude wing motion for flight. The time required for warm-up puts such insects at a considerable risk if they need to rapidly escape from predators. Here, we show that upon experiencing a sudden air-puff on the head, hawkmoths rapidly initiate flight bypassing the pre-flight shivering phase. This response is mediated by mechanosensory cephalic bristles that are buried under the scales on their head. Cephalic bristle mediated flight entails a stereotypic triggering of various flight-related reflexes including antennal positioning, foreleg extension, wing movement, and abdominal flexion. Some mechanosensory neurons underlying cephalic bristles arborize in the subesophageal zone (SEZ) and antennal motor and mechanonsensory center (AMMC), whereas most arborize in pro-, meso- and meta-thoracic ganglia which contain the motor circuitry for foreleg motion, flight, and abdominal flexion. Thermal recordings revealed that large-amplitude wing motion following cephalic bristle-stimulation occurs at lower thoracic temperatures than required for voluntary flight. Electromyogram recordings from steering and indirect flight muscles show significant variability in activation latency in response to cephalic bristle stimulus. The range of latency values among different muscles overlaps, suggesting that cephalic bristle stimulation activates steering muscles, thereby generating high-amplitude wing movement at lower thoracic temperatures. Concomitant activation of the indirect flight muscles initiates thoracic warm-up in preparation for longer flight. Thus, akin to locusts, the cephalic bristle system in hawkmoths rapidly triggers flight upon sensing danger, ensuring swift escape from potential threats.

## Introduction

The decision to initiate flight plays a pivotal role in the behavior of insects in various situations including evading predators, searching for food, finding mates, and defending territories. In some cases, flight initiation is voluntary and elicited by the internal triggering of flight-related actions (Taylor and Jones,1969; Lehane and Schofield 1981). Alternatively, flight-related reflexes may be triggered when a specific sensory cue, such as a sudden gust of air or a looming visual stimulus, signaling an approaching predator, surpasses a threshold value (e.g. Trimarchi and Schniederman, 1995). In locusts for example, flight can be initiated by gently blowing on the head region, thereby stimulating the cephalic wind-sensitive bristles (Weis-Fogh, 1949; Bacon, 1979). In *Drosophila melanogaster* and related flies, a sudden air current activates antennal mechanosensory afferents, which then initiate flight motor circuits, leading to flight (Strausfeld et al., 2014; Sadaf and Hasan, 2014; Sadaf et al., 2015). Alternatively, a looming visual stimulus can also quickly induce flight (e.g., Card and Dickinson, 2008).

Although most insects can initiate flight immediately after encountering such stimuli, certain larger insects including hawkmoths, butterflies, and bumblebees are slowed down by the need for their flight muscles to be in a physiologically prepared state to generate the requisite power to move their wings through their full amplitude (Stevenson and Josephson, 1990; Stevenson 1981). For instance, in hawkmoths, a warm-up routine is essential to bring their flight muscles into a prepared state. At thoracic temperatures below ca. 35° C, their flight muscles undergo twitches of a broad duration, thus causing the contractions of their antagonistic (i.e. wing-elevator and wing-depressor) muscles to overlap in time (Kammer, 1968). The co-contraction of antagonistic muscles at low temperatures means that the wings cannot go through their full excursion as required for flight, but instead exhibit rapid short amplitude flapping behavior called *pre-flight shivering* (Dorset, 1962; Kammer, 1971). This muscle activity generates heat, thereby elevating thoracic temperatures. During *pre-flight shivering* in hawkmoths, the frequency of wing vibration increases linearly with the temperature of the thorax (Heinrich, 1971). As the muscles warm up, their twitches become briefer in duration. Eventually, the antagonistic flight muscles twitch out-of-phase with one another, and the wings can now undergo their full excursion, thereby generating sufficient aerodynamic forces to fly. After a brief period of shivering, the flight muscle temperature remains elevated throughout the flight duration, thus allowing the insects to fly for longer durations (Dotterwich, 1928; Kammer, 1968). In the hawkmoth *Manduca sexta*, thoracic temperatures must be elevated to as high as 40°C to generate flight (Dorsett, 1962; Kammer 1971), and their maximum mechanical power output is substantially enhanced at higher temperatures in the range of 35-40°C, and cycle frequencies of 28-32 Hz, at which they voluntarily take flight or hover (Stevenson and Josephson, 1990).

In insects that require a pre-flight warmup, the long duration between flight initiation and take-off means that they are vulnerable to a sudden predatory strike from birds or bats. We know very little about sensory adaptations that allow such insects to rapidly initiate flight, overriding the need for pre-flight shivering behavior. Here, we show that a set of mechanosensory bristles on the head (henceforth called *cephalic bristles*) of hawkmoths, when stimulated by touch or a pulse of air, instantly elicit flight bypassing pre-flight shivering. These bristles are buried within a dense cover of scales in Lepidopteran insects and possess the distinct morphology of mechanosensory *sensilla chaetica* in insects. Our neuroanatomical investigations reveal that the mechanosensory neurons underlying these bristles project *via* long afferents into various parts of the brain and thoracic ganglia. Our behavioral and electrophysiological investigations on tethered moths show that, when stimulated with a sharp gust of air, these bristles activate various flight-related reflexes across the body. The wing movement is initiated at substantially lower thoracic temperatures than in hawkmoths which undergo voluntary flight. Together, these data reveal the fascinating sensorimotor system underlying the behavioral coordination of flight-related reflexes which enable rapid initiation of flight, bypassing the need for warm-up behavior.

## Methods

### Moth breeding

Oleander hawkmoths *Daphnis nerii* (Linnaeus, 1758) were bred under conditions of controlled temperature and humidity at the greenhouse facility on the National Centre for Biological Sciences campus in Bangalore, India. Moth larvae were reared on freshly cut leaves of *Nerium oleander L.,* which is their natural diet, and allowed to pupate in a netted container filled with sawdust. Post-emergence, adult moths were fed with approximately 3% sugar solution *ad libitum* before experiments. In the behavioral and neuroanatomical experiments described here, we used moths that were at least 1-day old. For the scanning electron microscopy experiments, we used 2-3 day old moths in which it was easier to remove the scales.

### Characterization of flight initiation mediated by cephalic bristles

#### High-speed videography

To tether the moths, we first descaled a small region on the dorsal surface of their scutum and affixed neodymium magnets of 3 mm diameter using Cyanoacrylate glue (Evobond^®^). These magnets were then attached to magnets of opposite polarity at the tip of a rigid metal rod, thus tethering the moth to the rod. We allowed the moths to adapt to dark conditions for at least 30 minutes before the beginning of each experiment. To stimulate the cephalic bristle region, we delivered air pulses of 250 ms duration through a solenoid valve connected to an air cylinder with a pressure regulator (Fig 1A). The opening and closing of the solenoid valve were controlled by a customized circuit. Three such trials were conducted for each moth with 45 minutes of inter trial interval.

**Figure 1:**
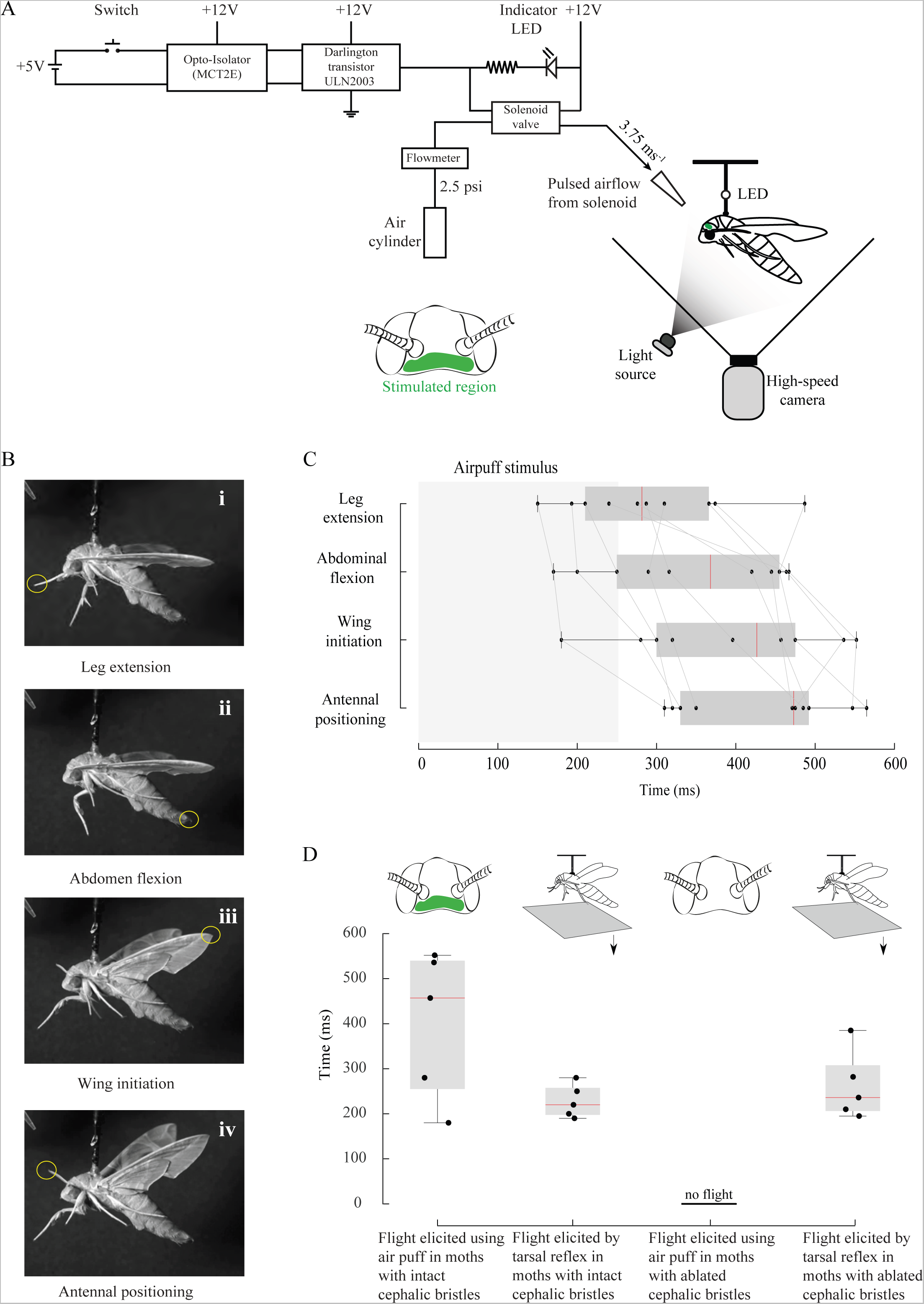
Response of moths to cephalic bristle stimulation by air pulse. (A) Schematic diagram of the experimental setup to deliver precisely timed air pulse stimulus of 3.75 m/s to the head of the moth. The region of the head that is stimulated is shown in the inset in green. (B) The position of tips of foreleg (B i), abdomen (B ii), wings (B iii), and antennae (B iv) post-stimulation. These points were tracked and digitized. (C) Behavioral response latency plot. Stimulus is delivered in the time duration t=0-250 ms (gray box). Response latencies are noted from the time instant of the first observable movement of foreleg, abdomen, wing, or antennae. In the box-and-whisker plots for each behavior, whiskers cover the entire range of the data, whereas boxes cover the lower and upper quartiles, and the median as a red line. All datapoints are overlaid. This convention is followed in all box-and-whisker plots in subsequent figures. A grey line connects the foreleg, abdomen, wing, and antennal movement latencies of each moth. (D) Box-and-whisker plots showing the response latency of flight initiation following air pulse stimulation for control moths with intact cephalic bristles and the corresponding tarsal reflex, followed by experimental moths with ablated cephalic bristles and the corresponding tarsal reflex. This figure shows that although air pulse mediated flight response is lost after cephalic bristle ablation, the moth can still be induced to fly using tarsal reflex.

The airspeed required to elicit flight was measured using a hotwire Mini anemometer (Kurz 490S, Kurz Instruments Inc., Monterey, CA, USA) in every trial. This allowed us to measure the actual airspeed delivered to the cephalic bristles, ranging from 1 to 6.5 m/s. The flight initiation responses of the moths were filmed from a lateral view at 3000 frames per second and an exposure time of 300 μs using a single high-speed camera (Phantom v7.1, Vision Research, Wayne, NJ, USA; Fig 1B i-iv). The wingbeat frequency of *Daphnis nerii* is about 30 Hz, and the frame rate we used was 3000 Hz, which is 100 times the wingbeat frequency. Thus, the frame rate captured the flight behavior with sufficiently high temporal resolution.

To determine the latency of the onset of movement of various appendages following stimulus delivery, we digitized various points on the moth’s body using the open-source Tracker software (Tracker-Video analysis and modeling tool, GNU general public license, version 3). The data were analyzed and plotted using MATLAB 2016a (Mathworks Inc., California; Fig 1C-D).

When the moth was immobile, the time series displaying the position of an appendage showed baseline noise, which is likely digitization error. We defined the onset of movement from the time instant when any change in position exceeded two standard deviations (i.e. 0.8mm) above the baseline noise. From this time instant, we next determined the behavioral latency as the time duration between the onset of stimulus and the onset of each behavior.

#### Mapping the cephalic regions sensitive to tactile stimulation of cephalic bristles

Unlike in locusts where the wind-sensitive cephalic bristles are localized in 5 symmetric fields (Weis-Fogh, 1949), the moth head is covered by scales and cephalic bristles which are not easily distinguishable. Hence, instead of stimulating individual bristles, we recorded behavioral responses to tactile stimulation of small regions on the head of a tethered moth (n=10). For these experiments, 0-to-1-day old healthy moths were cold anesthetized by placing them at −23°C, and tethered to a metallic rod using magnets. The moths were then allowed to rest for 45 minutes post-anesthesia. To test for recovery from anesthesia, we elicited flight using tarsal reflex and only active moths were used for experimentation. Tethered moths were positioned under a microscope with a camera (Nikon stereo microscope SMZ25, Nikon Instruments Inc., USA) to view the cephalic region at high resolution for precise stimulation. Whole-body response of the moths was captured with a web camera (Logitech C270 HD). A grid was overlaid on the image of the head in the software application associated with the microscope (NIS-elements D, version 4), and used as the reference to stimulate different regions across trials. A needle mounted on a manipulator (MM-3, Narishige, Japan) provided tactile stimuli to cephalic bristles. It was positioned close to the head of the moth and was translated gradually to stimulate randomly picked squares. The movements of antennae, wings, legs, and abdomen, and elicitation of flight were captured using the stereo microscope camera and the web camera. If this stimulus elicited flight, the moths were allowed to rest for 30-45 minutes, else they rested for 5 minutes between successive stimuli. The videos were manually analyzed and the response to each stimulated region was noted as the presence or an absence of an elicitation of flight, and movements of antennae, legs, wings, and abdomen, respectively. This allowed us to map the receptive areas of the moth head (Supplementary figure 1).

#### Cephalic bristle ablation

We ablated cephalic bristles to test if they were necessary for initiating flight upon air pulse stimulation. We systematically ablated all the scales and bristles in the cephalic bristle region using a toothbrush and followed the same protocol as in the cephalic bristle intact flight initiation experiment.

#### Analysis and statistical tests

A one-sample Kolmogorov-Smirnov test was used to assess the normality of our data. Because the null hypothesis -that our data followed a Gaussian distribution - was rejected (p ≤0.05), we used a non-parametric Wilcoxon rank sum test to compare the independent datasets that were not normally distributed. These tests allowed us to determine if the two datasets were derived from the same population.

### Thermal imaging of moth thorax during flight initiation

1-day old moths were affixed with a magnet on their ventral surface between the meso- and meta-thoracic legs. The moths were allowed to recover from this treatment for 30 minutes during which they were free to move around with the affixed magnet. To tether the moths, we used a ventral tether rod equipped with a magnet of opposite polarity and allowed it to rest for an additional period of 15 minutes. The temperature of the room was maintained at 25°C (starting temperature in all trials). To elicit flight *via* cephalic bristle field stimulation, we delivered a strong air current directed to the dorso-posterior (or crown) region of their head using the solenoid valve setup (Fig 1A, Fig 2A), and recorded the movements associated with flight initiation at 600 fps using a high-speed camera (Vision Research, v7.1). These movements were recorded from the front with the camera placed roughly orthogonal to the plane of wing movement (Fig 2A). This allowed us to determine the wing amplitude by quantifying the projected wing angle. To record voluntary flights, the moths were allowed to initiate flight on their own, which usually required us to wait between 5-20 minutes post tethering. The flights were usually preceded by a warm-up phase (Fig 2B-G). Simultaneously, all flight responses were filmed at 60 fps using a thermal camera (VarioCam HD 680, Infratech, Germany) which provided a dorsal view of the entire thorax. We ensured that the thorax of the moth was at room temperature (25°C) before starting the trials.

**Figure 2:**
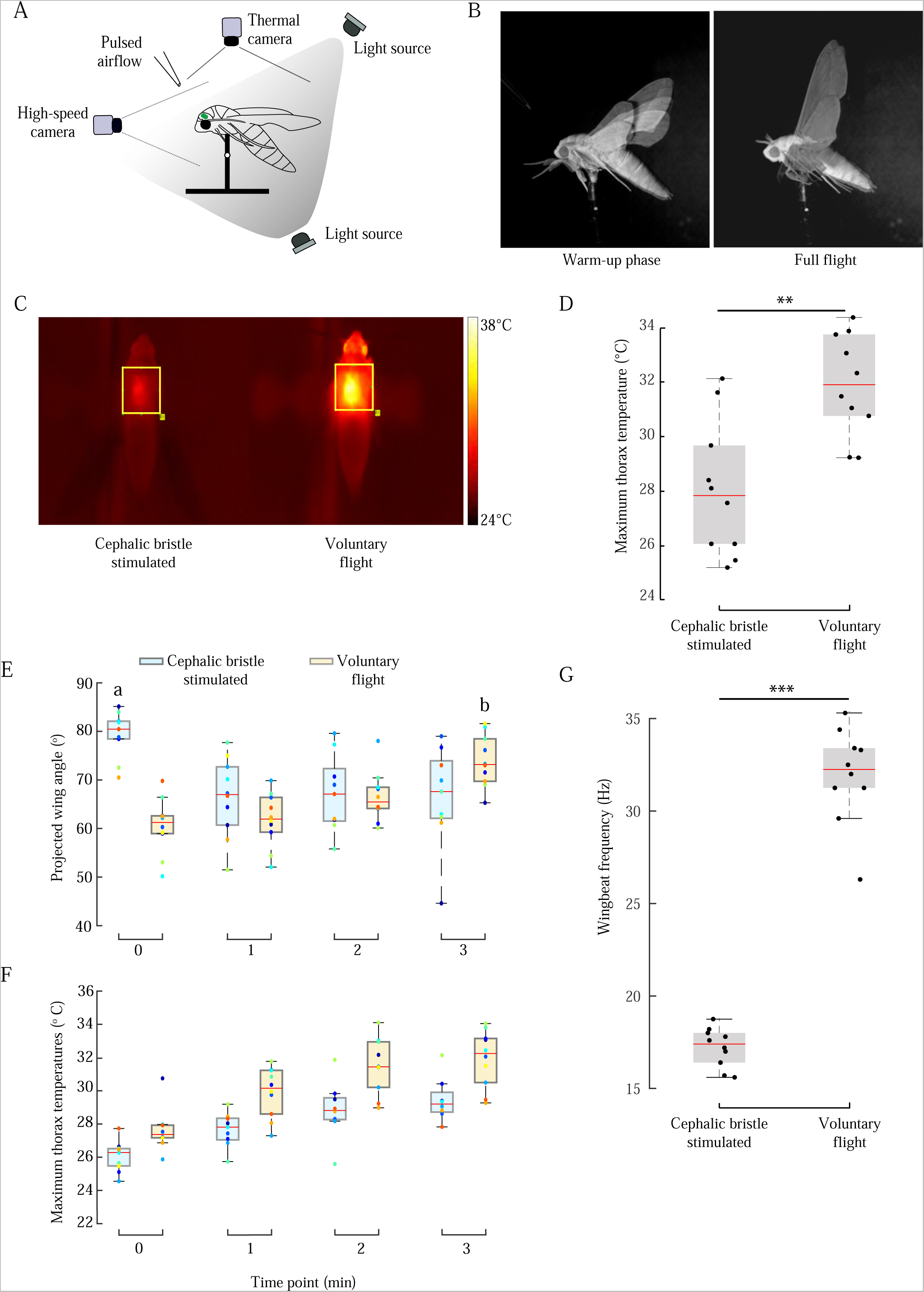
Thermograms of moth thorax. (A) Schematic diagram of the experimental setup to record the temperature of the moth thorax during voluntary and cephalic bristle mediated flight, using a dorsally placed IR camera. The moth was stimulated as described in the legend of Figure 1 A. (B) Snapshots of wing positions during pre-flight shivering phase during which stroke amplitudes are small, and during full flight when stroke amplitudes are large. (C) Thermogram showing the maximum temperature of the thorax during cephalic bristle mediated flight (left) and voluntary flight (right). Region of Interest used for measurement (thorax) is indicated by the yellow box. Temperatures can be read using the adjacent color bar. (D) Comparison of maximum thorax temperatures of moths during voluntary vs cephalic bristle mediated flight. ** indicates p < 0.01 (p = 0.004; Wilcoxon rank sum test). ‘ (E) Projected wing amplitudes of moths after cephalic bristle stimulation (blue box) and voluntary flight (orange) compared at 4 time points (0th to 3rd minute). Comparison indicated by boxes marked *a* vs *b* * indicates p<0.05 (p = 0.03; Wilcoxon rank sum test compared the wing amplitudes between time points where highest wing amplitude was recorded (time point 0 of cephalic bristle stimulated flight bout (box marked *a*) *vs* time point 3 of voluntary flight bout (box marked *b*)). (F) Maximum thorax temperatures of moths after cephalic bristle stimulation (blue box) and voluntary flight (orange box) compared across the same four time-bins as (E). Full amplitude flight is achieved by cephalic bristle stimulation without a significant increase in thorax muscle temperature. (G) Comparison of wing beat frequencies in moths with cephalic bristle stimulated using air pulse vs voluntary flight. *** indicates p << 0.005 (p= 0.00018; Wilcoxon rank sum test).

To synchronize both cameras, we fired a brief spark (generated by bringing a steel wool in contact with a capacitor) which was visible to both the high-speed and thermal cameras. This allowed us to measure the wing amplitudes while also measuring corresponding temperatures of the thorax. To verify the internal calibration of the thermal camera, we measured the temperature of boiling water using a clinical thermometer (Hicks Oval thermometer, Hicks Thermometers, Aligarh, India) while simultaneously filming it with the thermal camera. For high-speed filming, cool IR LED lights were used which did not register a rise in temperature by the thermal camera and did not heat the tethered moth. The videos were digitized using the DLTdv8 digitization tool (Hedrick, 2008).

### Scanning electron microscopy (SEM) of cephalic bristles

In Lepidoptera, the cephalic bristles are interspersed with scales making it difficult to identify their precise locations. Hence, for the scanning electron microscopy imaging of the bristles, it was essential to remove only the scales while keeping the bristles intact. In slightly older (3-4 days old) moths, scales can be removed more easily than in younger moths. Hence, we partially descaled the heads of such moths using a jet of air directed at their head. This method ensured the removal of the shallow-rooted scales, while retaining the mechanosensory bristles which are rooted more deeply (Fig 3A-C). The heads were then fixed in 4% paraformaldehyde for 8 hours, and gradually dehydrated in an ethanol series of 10-100% in 10% increments. The samples were dried using a critical point drying method (Leica EM CPD 300 Critical Point Dryer, Leica Microsystems, Wetzlar, Germany) to prevent distortions of the surface structures. Following this, we gold-coated the samples in a sputter coater (10nm, Emitech K550X, Quorum Technologies Ltd, West Sussex, UK) and imaged them in Field Emission Scanning Electron Microscope (Zeiss Merlin Compact VP/Zeiss Gemini SEM, Germany).

**Figure 3:**
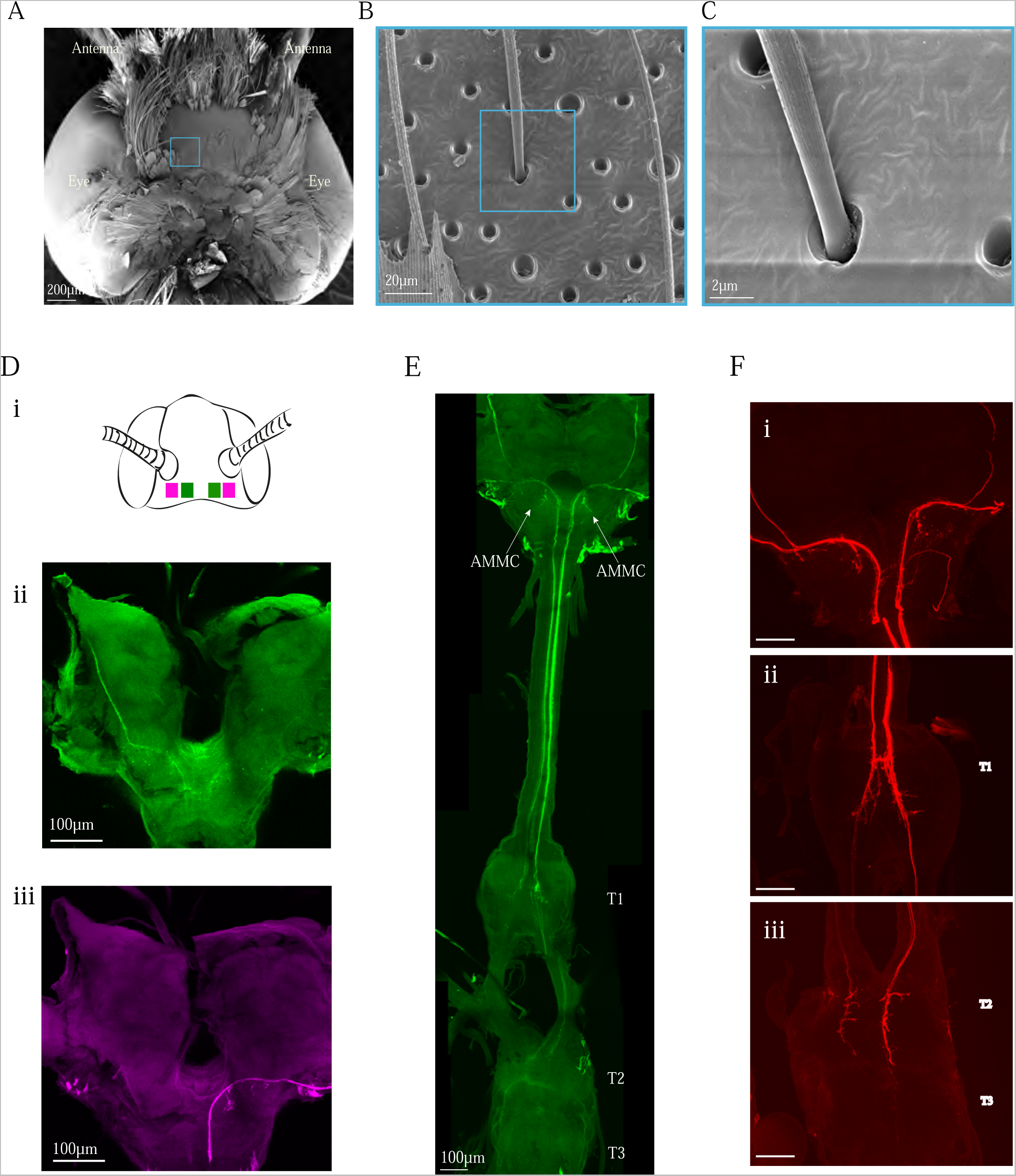
Morphology and neuroanatomy of cephalic bristles. (A) Scanning electron microscope (SEM) image of the dorsal view of head, with some scales removed to make the cephalic bristles visible. Blue boxes mark the region magnified in the subsequent images. Scale bar = 200 μm (B) Sparsely populated region showing scales (arrows) and cephalic bristles (blue box). Scale bar = 20 μm (C) Magnified view of the cephalic bristle. Scale bar = 2 μm (D) (i) Schematic of the head showing the regions targeted for fluorescent labelling of cephalic bristle sensory neurons. The primary sensory neurons of cephalic mechanosensors project to the brain *via* anterior (in green, ii) and posterior (in magenta, iii) tegumentary nerves. E) Ventral view of the brain and ventral nerve cord, showing a complete fluorescent labeled fill of the axonal projections from cephalic bristle neurons (green, Alexa Fluor 488). The three thoracic ganglia are marked T1 (pro-thoracic ganglia), T2 (mesothoracic ganglia) and T3 (meta thoracic ganglia). Scale bar = 100 μm. F) 2D projections of sensory neurons from the cephalic bristles (red) in the (i) subesophagal zone (SEZ) near the Antennal motor and mechanosensory centre (AMMC), (ii) prothorax, and (iii) meso and metathorax. Because the preparation involves keeping the moth alive for several hours after filling, the neuronal termini begin to disintegrate which causes the projections in the metathorax to always be somewhat blebby and faded. Scale bar = 100 μm.

### Fluorescent labeling of cephalic mechanosensory afferents

To label the cephalic mechanosensors, we first cold-anesthetized 1-day old hawkmoths by placing them in −23°C for 5-8 mins. The moths were then enclosed in a plastic syringe tube and immobilized using dental wax, with their dorsal side up. The scales and bristles in the cephalic region were completely removed. The cephalic bristle field region was dabbed with the fluorescent dye Alexa Fluor 488 (10,000 MW, Anionic, emission at 519 nm and excitation at 495 nm, D22910, Thermo Fisher Scientific) (Sant and Sane, 2018). These dyes are cytoplasmic markers, and hence require active transport of the cytoplasm to spread throughout the neuron.

Hence, the insect must be kept alive for some time after dye fill to allow its penetration through the whole cell. On the other hand, because the neuron is injured the distal processes begin to degenerate, which gives a blebby appearance to their projections. For neuronal dye fills, we must therefore contend with the trade-off between keeping the sample alive long enough to enable marking of the entire neuron, but not so long as to cause the distal ends to become blebs. Thus, the duration of this fill required some standardization. We fed the moths at regular intervals with sugar water for the duration they were to be kept alive. To dye-fill the projections of cephalic bristles in *Daphnis nerii*, we kept the moths alive in a hydrated chamber for 76-78 hrs. The specimens were fixed using 4% paraformaldehyde for 12-14 hours. After dissection, the brain was serially dehydrated in 50%, 60%, 75%, 90% and twice in 100% ethanol. It was then cleared with and mounted in methyl salicylate and imaged under Olympus FV1000 confocal microscope (Central Imaging and Flow Cytometry Facility (CIFF), NCBS, Bangalore, India) at 10-60 X magnification (Fig 3 D-F).

### Electromyogram recordings from flight muscles

#### Preparation

To determine the activity in the various flight muscles following cephalic bristle stimulation, we obtained electromyogram recordings from the steering and indirect muscles. The procedure for this is outlined below. Newly emerged (i.e. 8-10 hrs post-eclosion) moths were cold-anesthetized by placing them at −23°C for 5-8 mins. Each moth was then immobilized with its ventral side-up in a customized plastic syringe tube, using dental wax. The thorax of the moth was descaled, and the legs were clipped to allow easy insertion of electrodes. The cut-end of the legs was sealed with dental wax to avoid desiccation. Twisted pairs of silver wires with bare gold wire tip (12.5 μm thick) were used as electrodes. The recording electrode was inserted into any one of 4 types of flight muscles (basalar, subalar, upper axillary steering muscles, or the dorso-longitudinal indirect flight muscles) through a tiny hole made in the cuticle. After the whole prep was turned dorsal side up for recording, the reference electrode was placed on the surface of the thorax in saline surrounded by a wall of petroleum jelly outside the recording region. This preparation allowed us to record from the direct and the indirect flight muscles (Fig 4A, ii-vi).

**Figure 4:**
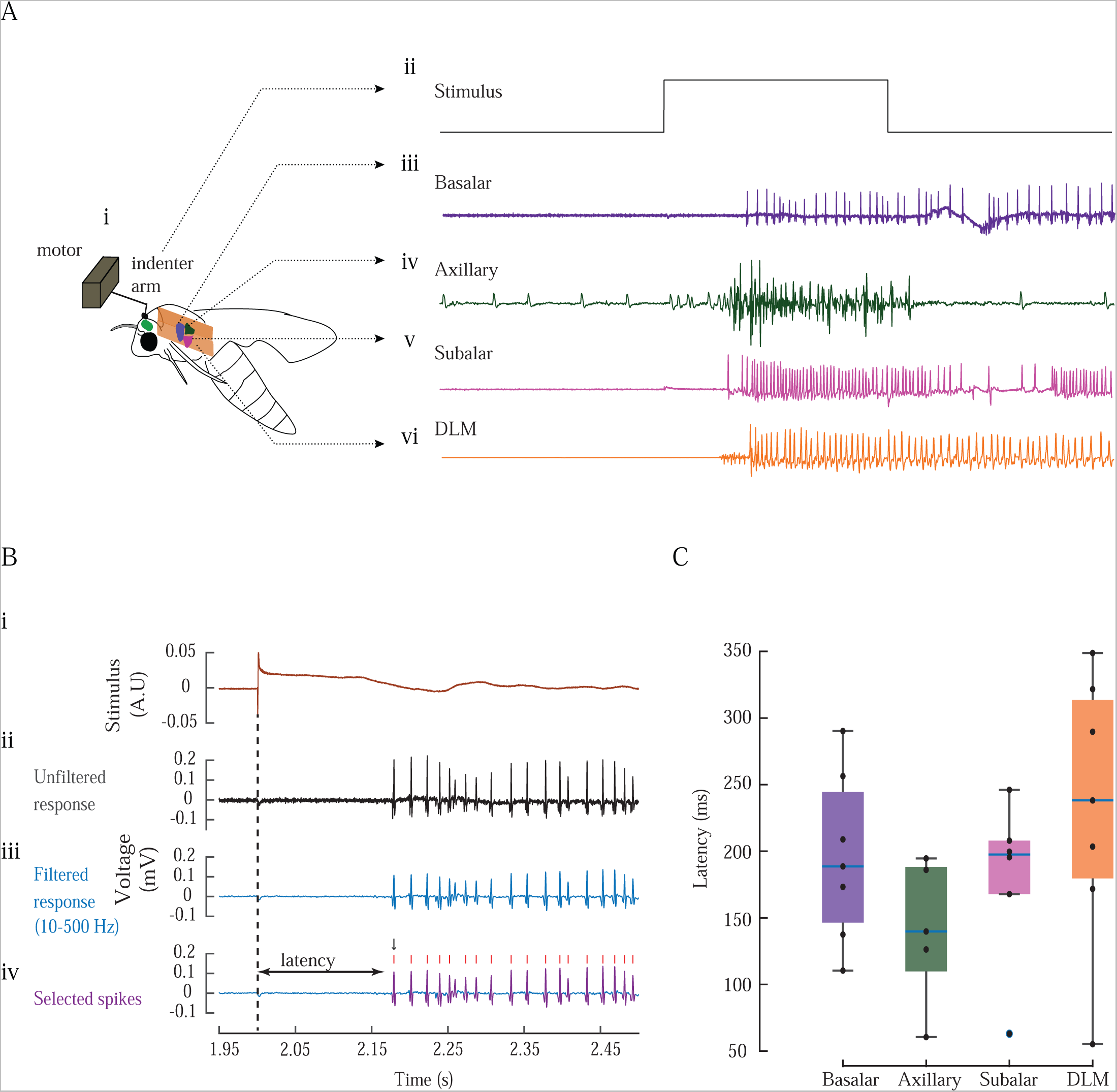
Wing muscles are activated simultaneously upon stimulation of cephalic bristles. (A i) Schematic diagram of the experimental setup for electromyogram (EMG) recordings from indirect flight muscles and steering muscles. (A ii) A stimulus pulse of 0.5 sec (black trace) is delivered by the motor arm. (A iii-vi) Response of the various wing muscles (basalar (iii), axillary (iv), subalar (v) and DLM (vi)) (B i-iv) Analysis pipeline for EMG data depicted using a representative recording from the basalar muscle as an example. All the trials analyzed are 2.5 sec long with 2 sec of baseline activity before stimulus and 0.5 seconds of stimulus duration. Here, the last 50 ms of baseline has been represented. (i) Feedback signal (in arbitrary units) from the servomotor upon contact with cephalic bristles and scales. (ii) Response recorded from basalar muscle (in mV) (iii) Bandpass filtered response (with 10-500Hz cut-off) (iv) Selected spikes after template matching (in purple) and the spike location (red ticks). Latency is measured as the time to first spike (arrow) after stimulus onset. C) Comparison of mean response latencies of 4 sets of flight muscles. Box-and-whisker plots indicate mean response latency for all trials of basalar (n=7 moths), axillary (n=5), subalar muscles (n=6) and DLM (n=7).

#### Stimulus

We stimulated the cephalic bristle in the crown region using a tactile stimulus delivered by a 3D printed circular disk of 1.67 mm attached to the arm of a moving coil motor (Mechanical stimulator 300C-I, Aurora scientific Ontario, Canada; Fig 4Ai). Upon delivering step voltage stimulus to the indenter arm, the circular disk attachment mechanically touched the scales and bristles in the crown region thus physically stimulating the mechanosensors (Fig 4A i-vi). After assessing the success of stimulation based on the feedback signal from the motor (Fig 4B i) and response from the corresponding muscle (Fig 4A iii-vi), we fine-tuned the position of the indenter arm for all the subsequent trials. Two kinds of stimulus protocols were provided: a single step stimulus with a duration of 0.5 seconds and three steps of the same duration with an interval of 4.5 seconds between each step. For each moth, three trials of single-step and three trials of three-step stimuli for each muscle was recorded. A pre-stimulation baseline activity was recorded for 2 seconds before every stimulus. The precise instant of the stimulus could be registered using the force feedback signal from the motor when a contact with the bristles was made. An interval of 25 minutes was given between trials to allow the moth to return to its undisturbed state.

#### Analysis

We recorded from 8 individuals with 6 trials on each of the 4 muscle groups (i.e. total of 192 trials; case example in Fig 4A i-vi). Of these, 110 trials were used for analyses (Axillary-5 individuals, 28 trials, Basalar-7 individuals, 25 trials, DLM (Dorso-longitudinal muscles) −7 individuals, 29 trials and Subalar-6 individuals, 28 trials). We discarded data if the recording had unexpected variability in stimulus feedback signal during the resting period, or if the muscle exhibited high spontaneous baseline activity that continued even after stimulation in which case the effect of stimulus was hard to discern. Such cases most likely reflected that the head was in continuous contact with the indenter arm, which meant continuous stimulation of the cephalic bristle. Cephalic bristle stimulation was strong enough to sometimes cause minor head movements despite immobilization of the neck joints.

#### Filtering and spike sorting

The raw data (Fig 4B i,ii) were bandpass-filtered using a 3rd order Butterworth filter with cut-offs at 10 Hz and 500 Hz (Fig 4B iii). In many cases, the response following the first step stimulus in a three-step protocol trial heavily overlapped with the occurrence of the next stimulus. To avoid ambiguity, a total length of 2.5 seconds of recording was used for all analyses which included the pre-stimulation period of 2 seconds and 0.5 second of stimulus duration. Spike detection was performed by a custom-written program in MATLAB employing a template matching algorithm. (https://github.com/chinmayeelm/EMG_spike_detection). Thresholds on the amplitude (>5mV) and width of the spike (2ms-11ms) were applied on the identified spikes to eliminate potential non-spikes (For details of the methodology, please refer to supplementary Methods). The number of templates used in this scheme depended on the variations in spike features. Redundant overlapping matches were eliminated. The first spike occurring after the onset of stimulus, with amplitude larger than baseline spikes, was used for latency measurements (Fig 4B iv). Pairwise comparison of latencies of different muscle groups showed no significant statistical difference (Fig 4C).

## Results

### Moths initiate flight in response to air pulse stimulation of their cephalic bristles

When a specific region of the head was stimulated with an air pulse, the hawk moths rapidly initiated flight. This response was composed of multiple actions including leg extension (latencies ranging from 152 ms - 564 ms post-stimulus-onset), and followed by abdomen flexion (176 ms - 467 ms), initiation of wing flapping (188 ms - 547 ms), and antennal positioning (318 ms - 648 ms) (Fig 1 B, C; n=10 moths, trials = 3 each). These responses largely occurred in a definite sequence; in 5 out of 10 individuals (20 out of 30 trials), the sequence began with the movement of prothoracic legs followed by abdominal flexion, wing movement and antennal positioning (Fig 1B i-iv; Supplementary Table). The flight was initiated typically at airspeeds above 3 m/s and the probability of flight initiation saturated at around 4.5 m/s (Supplementary Fig 2). At airspeeds below 3m/s and above 4.5 m/s, leg and abdominal movements were sometimes observed along with low-amplitude wing flapping but not flight initiation, suggesting that the response had saturated.

These data indicated the possibility of mechanosensory structures on the head which act as airflow sensors, similar to other insects (e.g. locusts, Weis-Fogh, 1949, Camhi, 1969). Because the moth’s head was densely covered with scales, it was initially difficult to identify specific mechanosensory bristles. To ascertain that mechanosensory bristles may be involved in flight initiation, we removed all scales and bristles on the head by brushing them off with a toothbrush. Any remaining bristle-like structures were removed with the help of forceps. When the head area was again stimulated with an air pulse, we did not observe any flight initiation response, even though the moth retained its ability to fly if stimulated by other means. For instance, in these moths, flight could be elicited upon sudden loss of tarsal contact, a phenomenon referred to as *tarsal reflex* (Fig 1D, n=10, 3 trials per moth) (Weis-Fogh, 1951). These data indicated the presence of mechanosensory cephalic bristles which, when stimulated by an air pulse of a magnitude greater than 3 m/s, elicit the flight response involving sequential activation of various flight-related actions. Importantly, none of the moths displayed pre-flight shivering behavior prior to flight initiation *via* cephalic bristle stimulation.

### Pre-flight shivering phase is bypassed during cephalic bristle-mediated flight initiation

For sustained flight over long durations, hawkmoths must warm up their flight muscles (Dorsett 1962, Kammer 1971, Heinrich and Bartholomew 1971). However, the requirement for such warm-up behavior means that moths cannot elicit high-amplitude wing flapping until their muscle temperature is high (37-41°C in *Manduca sexta*, Heinrich, 1971, 34-45°C in *Deilephila nerii (*Dorsett, 1962; same as *Daphnis nerii* by Hübner, 1819).

The rapid elicitation of flight following cephalic bristle stimulation led us to the hypothesis that the standard warm-up process is bypassed when cephalic bristles are stimulated, eliciting flight with full-amplitude wing movement. To test this hypothesis, we compared the thoracic temperatures of moths in which flight was voluntary, with moths in which flight was elicited *via* cephalic bristle stimulation. In these experiments, we recorded the thoracic temperatures of the moths using an infrared-sensitive thermal camera, while simultaneously recording their wing movements with a high-speed camera (Fig 2A-B). In moths subjected to air-pulse stimulation on the head, high-amplitude wing flapping occurred at thoracic temperatures significantly lower than those necessary for self-initiated flight (Fig 2B-C). Specifically, in *Daphnis nerii*, flight due to cephalic bristle stimulation occurred at thoracic temperature of only about 28 °C (range 25-32 °C, Fig 2C-D) whereas during voluntary flight the thoracic temperatures were as much as 32 °C (range 29-35 °C, Fig 2D; see also Dorsett 1962; 40°C for voluntary flight in *Manduca sexta*, Kammer 1971).

We measured the stroke amplitude and corresponding thoracic temperatures in 1-minute intervals immediately following cephalic bristle stimulation or flight onset in the case of voluntary flight, for a total of 3 consecutive minutes. Following cephalic bristle stimulation, the stroke amplitude ranged between 72-85° (median= 80.4° in the first minute; blue box, Fig 2E) at thorax temperatures ranging from 25-27°C (Fig 2F), but in the subsequent time intervals, the amplitude fell to a range of 44-79° (median = 67.6° in the third minute; Fig 2E; thorax temperature 27-32° C; Fig 2F) at which it remained steady. In contrast, the stroke amplitude of the moth which voluntarily initiated flight was initially at 50-69° (median= 61.2° in the first minute; yellow box, Fig 2E), but steadily increased to about range 65-81° (median=73.3° in the third minute; yellow box; Fig 2E) after the thorax warmed up to 29-34 °C. Thus, cephalic bristle stimulation elicited high amplitude flapping at low thoracic temperatures. After the initial burst of wing flapping following cephalic bristle stimulation, the moth settled into a low-amplitude mode of flapping which is typical of warm-up phase of voluntary flight. Wing amplitudes in the first few seconds of cephalic-bristle mediated flight were slightly greater than those achieved in a flight post-warmup. The wingbeat frequencies of moths flying after air pulse stimulation of their cephalic bristles was significantly lower than moths flying voluntarily (Fig 2G; Wilcoxon rank sum test, p=0.00018). These observations indicated that the initial burst of flight activity caused by cephalic bristle stimulation bypasses the standard warm-up routine. However, such flight cannot be sustained over a long period as the thoracic temperatures need to be elevated for sustained flight over long durations.

### Cephalic bristle mechanosensory afferents project into the brain and thoracic ganglia

In the behavioral data described above, the rapid elicitation of flight following the mechanical stimulation of cephalic bristles suggested the hypothesis that these bristles directly activate flight-related reflexes. Testing this hypothesis required us to first identify the structure of these bristles, which are covered by the scales on the head. We first identified specific areas on the head where a mechanical (tactile) stimulus caused flight initiation (Supplementary Fig 1). After heavily descaling the crown region (see methods), we imaged it using a Scanning Electron Microscope (SEM) and identified bristles in this region that resembled the typical structure of mechanosensory bristles (Fig 3A-C). The bristles described here have grooved bristle shafts embedded in a socket of about 2 μm diameter with deep roots, which are features typical of *sensilla chaetica* and are very distinct from the scales (Fig 3A-C, e.g. Field and Matheson, 1998, Snodgrass, 1935).

We next removed both the scales and bristles from this region of the head and dabbed this shaved area with the fluorophore dye Rhodamine dextran or Alexa fluor 488 to label the underlying sensory neurons (Fig 3D). The primary axons of the neurons spanned the entire distance from the head to the thoracic ganglion. The axons entered the brain *via* a pair of anterior and posterior tegmental nerves (Fig 3Di-iii) and passed through the cervical connective into the thoracic ganglia (Fig 3E). Along the way, these neurons arborized in the subesophagal zone (SEZ) and antennal motor and mechanosensory center (AMMC) (Fig 3F i), and further down in the pro-thoracic (T1) ganglion (Fig 3F ii) and meso- and meta-thoracic ganglia (Fig 3F iii), where the flight motor neurons for the front and hind wings are located (Rind, 1983).

### Flight muscles are activated in response to stimulation of cephalic bristles

Under normal conditions, large-amplitude wing movements require thoracic temperatures above 32°C which are generated by preflight shivering. Thus, for a voluntary flight bout, there is a time-lag between initiation of thoracic warmup and flight initiation, which can pose a severe disadvantage in case the moths need to rapidly initiate flight. Unlike voluntary flight, the cephalic bristle-mediated flight bypasses the warm-up process, allowing the moths to rapidly initiate high-amplitude wing flapping even when thoracic temperatures are lower. We proposed the hypothesis cephalic bristle stimulation activates steering muscles which rapidly enhance wing amplitude. To test this hypothesis, we provided tactile stimulus to the cephalic bristles using a moving coil motor (Fig 4A i,ii), and recorded electromyograms from diverse flight muscles including the basalar (Fig 4Aiii), axillary (Fig 4Aiv), and subalar (steering muscles) muscles (Fig 4Av), and also the DLMs (indirect flight muscles; Fig 4Avi). Using tactile stimulus rather than an air pulse allowed us to better characterize the time instant of stimulation, for latency estimation. The axillary muscles typically displayed baseline spontaneous activity before the stimulus, in contrast to the DLM, subalar, and basalar muscles which showed no such pre-stimulus activity (Fig 4Aiii-vi). In all cases, there was a clear burst in activity in the indirect and steering muscles after the onset of the stimulus to the cephalic bristles. The recordings were bandpass filtered and spike sorted, to determine latency of the response (Fig 4Bi-iv). The latencies of these responses were highly variable with their mean latencies ranging from 110-290 ms in basalar muscles (25 trials in 7 individuals), 60-194 ms in axillary muscles (28 trials in 5 individuals), 168-246 ms in the subalar muscles (28 trials in 6 individuals) and 55-348 ms in DLMs (29 trials in 7 individuals). The lowest values of latency still exceed 50 ms, which makes it unlikely that cephalic bristle afferents directly synapse on to the motor neurons of the steering or indirect flight muscles, even though the activation can sometimes be within timescales of a few wing beats. Due to the high variability in these latency measurements, a clear sequence of muscle activation could not be established within the flight muscles. We did not observe statistically significant differences among the groups (p=0.224; Kruskal-Wallis test). Pairwise comparisons among the muscle groups using the Wilcoxon rank sum test also yielded p-values of 0.267 (axillary-basalar), 0.125 (axillary-subalar), 0.106 (axillary-DLM), 0.835 (basalar-subalar), 0.455 (basalar-DLM), 0.294 (subalar-DLM).

## Discussion

The behavioral experiments presented here show that air pulse stimulus to cephalic region activates a coordinated set of flight reflexes, including antennal positioning, wing initiation, leg extension, and abdominal flexion. Thermal imaging of the moth thorax shows that such rapid flight activation *via* cephalic bristle stimulation bypasses the standard warm-up routine, thereby allowing high-amplitude wing motion at lower thoracic temperatures. The anatomical investigations point to a set of mechanosensory bristles interspersed with scales on the head of the moths, mediating this behavior. Ablation of these bristles abolishes the air-pulse stimulated flight-initiation. The neuroanatomical investigation shows that the axonal tracts of the mechanosensory neurons underlying the cephalic bristles pass through the brain, the cervical connective, and project to the meso- and meta-thoracic ganglia which contain the flight motor neurons. Finally, the electrophysiological data show that the stimulation of cephalic bristles activates both the steering and indirect flight muscles *albeit* with variable and long latencies suggesting the presence of intermediate interneurons between cephalic bristle afferents and flight muscle motor neurons.

### A comparative view of air pulse mediated flight initiation in insects

Air pulse mediated flight initiation and maintenance has been studied in diverse Pterygote insects ranging from Palaeopteran insects such as dragonflies (Sveshnkov, 1973) to Neopteran groups such as the Orthopteran locusts (Weis-Fogh, 1949; Camhi, 1969), Hymenopteran honeybees (Khurana and Sane, 2016), Lepidopteran moths (Sane et al., 2007; Sane et al., 2010) and Dipteran flies (Mamiya et al., 2011, Sadaf et al., 2012). In these insects, antennal mechanosensors, principally the Johnston’s organ, plays a key role in detecting air flow. JO afferents typically project into the antennal motor and mechanosensory centre (AMMC) (e.g. Krishnan and Sane, 2015) from where descending interneurons may convey this information to the flight circuits.

Compared to antennae, the role of cephalic bristles in the context of air flow detection and its subsequent role in flight control has been explored in fewer insects. This system is best studied in locusts, where cephalic bristles were first described by Weis-Fogh (1949) and subsequent studies explored their role in the control and maintenance of wing-beat frequency and phase in flying locusts (Camhi, 1969; Weis-Fogh, 1956). A series of studies followed these initial results, yielding a detailed picture of the underlying circuitry and the role of cephalic bristles in flight via the stimulation of the tritocerebral commissure giant (TCG) interneuron which integrates visual and mechanosensory inputs (Bacon and Mohl, 1983). These studies revealed that, like cercal bristles in cockroaches and crickets (Camhi and Tom, 1978, Steinmann et al, 2006; Dangles et al, 2006, Casas et al, 2010), the cephalic bristles signal the presence of a sudden gust of wind and cause flight initiation, and thus may be involved in generating escape responses.

Besides locusts, cephalic bristles have been reported in the hematophagous bug *Triatoma infestans,* in which central projections of the bristles were mapped using cobalt fills (Insausti and Lazzari 2000) but not much is known about their effect on behavior in general and flight in particular. Similar to the projection patterns in locusts (Tryer, Bacon and Davies, 1979) and moths in the current study, the cephalic bristle axons in these bugs are carried by the anterior and posterior tegumentary nerves into the brain. The axons project to the meso- and meta-thoracic ganglia *via* the neck connective, with branches in the SEZ and AMMC and the prothoracic ganglia. Although their role in locust flight is well-documented, the function of cephalic bristles in *Triatoma infestans* is not well-studied. Notably, although *Triatoma infestans* can fly, they primarily walk and hence the cephalic bristles may play a role in anemotactic orientation towards odor sources. Because these two insects are phylogenetically distant, the cephalic bristle system may be a conserved feature across insects (Insausti and Lazzari 2000).

The data presented in this paper indicates that a cephalic bristle system in *Daphnis nerii* is very similar and perhaps even homologous to Orthopteran locusts and Hemipteran bugs. There are several points of similarity between locusts and moths. First, as reported in locusts, air stimulus to the cephalic bristles causes rapid flight initiation in moths (Weis Fogh 1949). Second, similar to locusts, the cephalic bristles are sensilla chaetica, a type of trichoid sensilla (Snodgrass, 1935). Third, as in the locust *Schistocerca gregaria*, the central projection patterns of the mechanosensory neurons project via tegumental nerves into the brain, and their central projection patterns are similar (Guthrie, 1964; Tryer, Bacon and Davies, 1979).

It was not possible in our study to clearly identify bristle fields in moths, due to the profusion of scales on the head. In locusts, a majority (75%) of these projections terminate in the SEZ, with only ∼20% projecting into the thoracic ganglia (Bacon and Tryer, 1979; Tryer, Bacon and Davies, 1979). The inputs that terminate in the SEZ feed into the Trictocerebral commissure giant (TCG) system, which integrates inputs from various bristle fields and other modalities including the antennal mechanosensory inputs and visual inputs (Bacon and Tryer, 1978, Bacon and Mohl, 1983, Mohl and Bacon 1983). In contrast to locusts, a majority of cephalic bristle afferents in moths appear to project directly into the thoracic ganglia, with some branches in the SEZ and AMMC (Fig 3E, F). Despite these projections, a TCG-like system cannot be ruled out in the case of moths as the neuroanatomical study described here was not designed to label interneurons. We were also unable to establish a directionality to the response for individual bristle responses, although the anatomy of the basal shaft of the bristles is asymmetric suggesting that at least some directional response may be conferred upon each bristle due to this asymmetry. Finally, from a functional viewpoint, even a brief stimulus to the cephalic region of a quiescent moth can elicit flight that can last for several minutes. Thus, there may be a strong neuromodulatory influence causing the flight to be sustained. In locusts, the excitatory role of octopamine has been well-established in flight (Orchard et al, 1993). Application of octopamine to the thoracic ganglia or stimulation of cephalic bristles releases flight-like rhythm at all developmental stages in locusts, although only the adults possess wings (Stevenson and Kutsch, 1987). Based on these data in locusts, we speculate that stimulation of the cephalic bristle in moths may also exercise a neuromodulatory influence to cause the release of flight rhythm. These neuromodulatory intermediates need to be explored in future studies.

### Flight initiation and the coordination of reflexes

In contrast to the voluntary flight behavior in *Daphnis nerii,* an air pulse stimulus to the cephalic bristles initiates flight rapidly, bypassing the warm-up routine and generates large amplitude wing motion *albeit* at lower thoracic temperature and wingbeat frequency (Fig 2D-G). EMG recordings from the various steering muscles and indirect flight muscles show the activation of all these muscles by cephalic bristle stimulation. Because temperature plays a significant role in muscle activation in moths (Kammer, 1971; Stevenson, 1990), locusts (Weis-Fogh, 1956) and bees (Sotavalta 1954), the observation that moths can initiate large amplitude flight at low thoracic temperatures is unexpected. Moreover, such flight is often transient, before switching to a warm-up routine.

We propose the following scenario to explain the above results. First, the mechanosensory neurons underlying the bristles innervate the thoracic ganglia, and activate local flight-related reflex circuits to rapidly jumpstart flight (Fig 5). However, the flight behavior thus activated cannot be sustained over long durations because the flight muscles are not sufficiently warm. Steering muscle activity thus plays a key role in generating a few large amplitude wing strokes eliciting a short flight bout. Simultaneously, the activation of the indirect flight muscles initiates a warm-up routine that prepares the moth for longer duration flight bouts. Second, the activation of cephalic bristles causes a systemic activation of various other flight-related reflexes including antennal positioning, head stabilization, wing motion, and abdominal flexion. These responses ensure that the moths can briskly take flight and escape from the vicinity of a predatory bat or bird. Having flown out of reach of the predator, they can perform a sustained warm-up routine to ensure long duration flight bouts.

**Figure 5:**
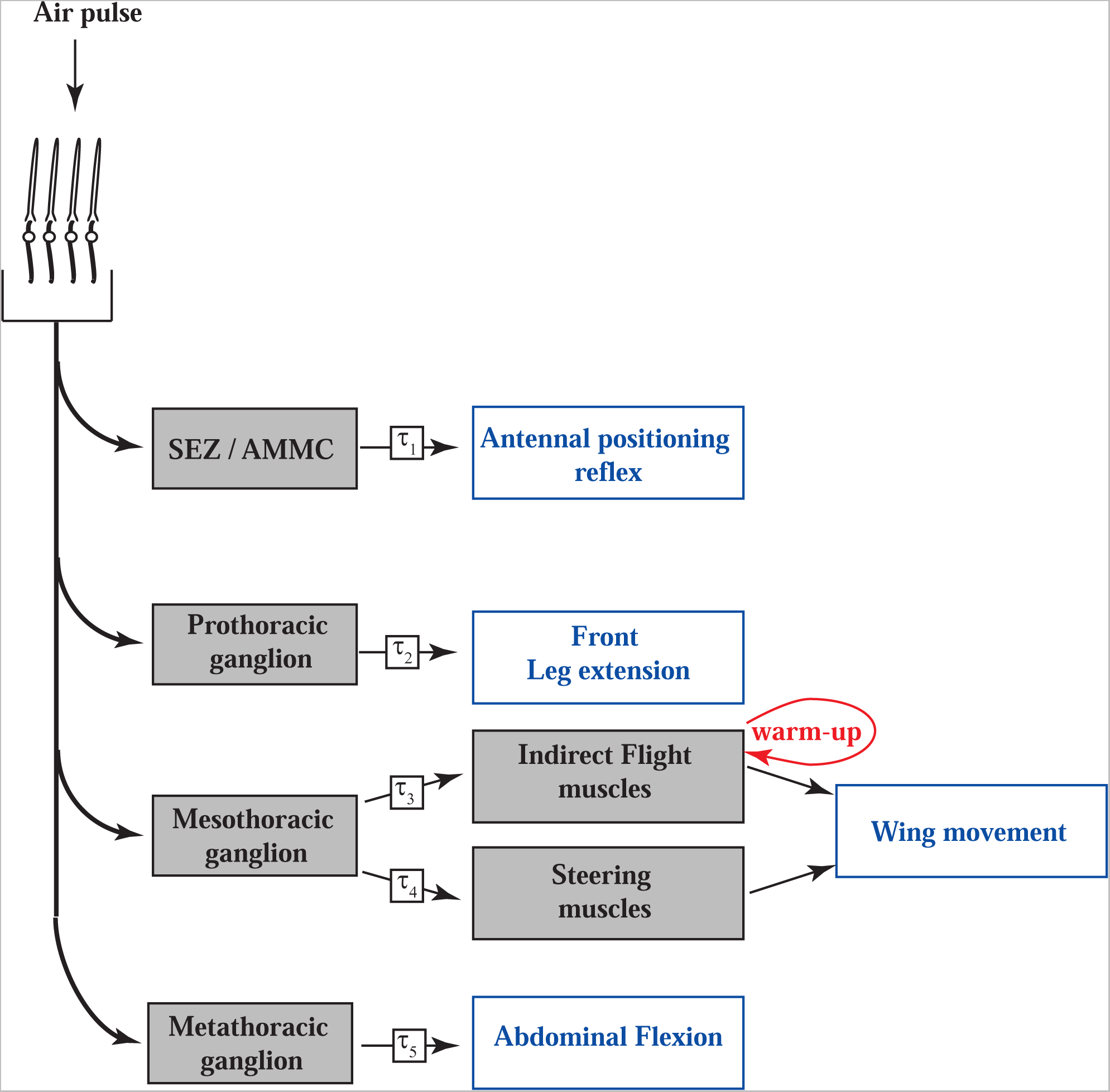
Mechanism of cephalic bristle mediated flight initiation. An air pulse to the cephalic bristles sequentially activates flight-related reflex loops which include antennal positioning, foreleg extension, IFM-driven and steering muscle-driven wing movements and abdominal flexion with latencies of τ_1_-τ_5_ respectively. According to this conceptual model, the activation of indirect flight muscles initiates a feed-forward warm-up process (in red) which helps elevate the thoracic temperatures in preparation for longer flight.

### Threshold values of cephalic bristle stimulation

In all the behavioral experiments described above, an air pulse stimulus greater than 3.0 m/s was required to elicit flight. Cephalic bristles are buried amidst a mass of scales, which act as a mechanical filter to lower their responsivity to air pulse stimuli, thereby increasing the threshold value of the air pulse stimulus required to elicit flight. This ensures rapid activation of flight only when the air pulse is sudden, as would occur when a predator is relatively close or if there is a sudden gust of wind. However, this heightened threshold may also influence other aspects of the flight performance in moths, such as long-distance migratory flight in Lepidoptera in which insects typically fly at speeds exceeding 3 m/s. For example, the neotropical moth *Urania fulgens* typically flies at speeds of 3.8 m/s (Sane et al 2010) whereas Caribbean butterflies *Phoebis sennae* fly at airspeeds ranging from 4.04 to 4.24 m/s (Srygley, 2001). At these speeds, we hypothesize that continual stimulation of the cephalic bristles helps maintain elevated levels of the excitatory neuromodulators, thus ensuring flight bouts of much longer duration than in non-migrating moths and butterflies. This constant airflow overlies a periodic signal that the cephalic bristles receive from the flapping wings (Sane 2006, Sane and Jacobson 2006), which may also serve to regulate wing motion. In locusts and other insects, similar mechanisms are known to regulate the phase and frequency of their wings following stimulation of cephalic bristles (Horsmann et al 1983). Lastly, the threshold for air pulse mediated flight stimulus is substantially higher when their legs are in contact with a substrate. Whereas dorsally tethered moths initiate flight at 3.0 m/s (Supplementary Fig 2), ventrally tethered moths in which tarsi were in contact with the tether, required air speeds of 4.5m/s and above. This suggests that tarsal contact inhibits flight and counteracts the excitatory effect of the cephalic bristle mediated elicitation of flight. This is similar to previous studies on wind sensitive bristle systems in locusts, in which flapping flight was activated only when legs were not in contact with the substratum (Weis-Fogh,1949).

## Conclusion

Together, these data show that stimulation of mechanosensory cephalic bristles causes rapid activation of flight muscles. This stimulus is transduced via long primary afferents that span past the cervical connective and arborize in the mesothorax, where they activate the flight motor circuit including both the indirect and the steering flight muscles. This ensures high amplitude wing movements that elicits short-duration flight bouts despite the lower temperature of the thoracic muscles. This is reminiscent of the escape behaviors in crickets and cockroaches following activation of the posterior cercal bristle. In most cases, after the rapid flight initiation, moths enter a warm-up mode which can then ensure more sustained flight bouts. We hypothesize, these sustained flight bouts point to excitatory neuromodulation of the flight circuits. Further studies are needed to address this hypothesis and to determine the key components of the cephalic bristle-to-flight neural circuit. Moreover, this circuit may be conserved in diverse and distantly related insects, including moths, locusts, and bugs. Further studies are required to fully explore and compare the structure and function of this circuit across insects.

## Acknowledgements and Dedication

During the course of our investigation, we learned that late Prof Edward Arbas from the University of Arizona had begun research on cephalic bristles in the hawkmoth *Manduca sexta* before his untimely passing in a tragic accident in 1996. Although his findings were never formally published, he had shared some results through a poster presentation at a Society of Neurosciences meeting. We dedicate this paper to the late Professor Ed Arbas to acknowledge his contributions to this field, and thank Prof Mark Willis for bringing his work to our attention. We thank the Central Imaging and Flow Cytometry (CIFF) and Scanning Electron Microscopy facilities at NCBS, Bengaluru, India, late Prof Chandrashekhara, GKVK for help with cephalic bristle morphometry, Megha Ramachandra for mapping cephalic bristle regions. We thank Dr. Shuchita Soman for her valuable insights and feedback on the manuscript. Funding was provided by Air Force Office of Scientific Research, Grant/Award Numbers: FA2386-11-1-4057, FA9550-16-1-0155; Department of Atomic Energy - Government of India, Grant/Award Number: 12-R&D-TFR-5.04-0800 to SPS.

## Supplementary Figure legends

**Supplementary Fig 1: Distribution of cephalic bristles on the head of *Daphnis nerii*** Responses of 10 moths to tactile stimulation of cephalic bristles overlaid on the schematic of a moth head. Regions on the moth head that elicited any movement of antennae, legs, wings, or abdomen, or elicited flight are marked by translucent squares. Darker areas, as seen in the grey scale, are the regions that elicited a behavior in more moths.

**Supplementary Fig 2: Threshold for flight initiation in cephalic bristle stimulated moths in the presence and absence of tarsal contact**

Flight is initiated when stimulated with an airspeed of 3 m/s or higher. The probability of flight initiation also reaches a maximum at stimulus air pulse speed of 4.5 m/s or higher, when the tarsi are not in contact with the substrate. However, when the tarsi are in contact, the stimulation threshold increases to 4 m4m/s when the tarsi is in contact. At airspeeds between 2.5 to 3 m/s, only prothoracic leg movements were observed.

**Supplementary Table: Sequence of activation of flight-related reflexes post stimulation of cephalic bristles**

Stimulation of cephalic bristles with an airspeed of 3.75m/s activated flight-related reflexes like leg extension, abdominal flexion, wing initiation and antennal positioning in a specific sequence. Color shades represent the order of occurrence, with darker shades indicating later occurrence in the sequence (n = 10 moths, 3 trials each).

## References

Bacon J., and Tyrer. M (1978). The tritocerebral commissure giant (TCG): A bimodal interneurone in the locust, *Schistocerca gregaria*. Journal of comparative physiology.

Bacon, J., and Tyrer, M. (1979). The innervation of the wind-sensitive head hairs of the locust, *Schistocerca gregaria*. Physiological Entomology, 4(4), 301–309.

Bacon, J., & Möhl, B. (1983). The tritocerebral commissure giant (TCG) wind-sensitive interneurone in the locust: I. Its activity in straight flight. Journal of comparative physiology, 150, 439–452.

Camhi, J. M. (1969). Locust wind receptors: III. Contribution to flight initiation and lift control. Journal of Experimental Biology, 50(2), 363–373.

Camhi, J. M., & Tom, W. (1978). The escape behavior of the cockroach Periplaneta americana: I. Turning response to wind puffs. Journal of comparative physiology, 128, 193–201.

Casas, J., Steinmann, T., & Krijnen, G. (2010). Why do insects have such a high density of flow-sensing hairs? Insights from the hydromechanics of biomimetic MEMS sensors. Journal of the Royal Society interface, 7(51), 1487–1495.

Card, G., & Dickinson, M. H. (2008). Visually mediated motor planning in the escape response of Drosophila. Current Biology, 18(17), 1300–1307.

Card G. M., (2012). Escape behaviour in insects. Current opinion in neurobiology. 22:180–186

Chapman RF (1982) The insects: structure and function. 3rd edition. Cambridge (Massachusetts): Harvard University Press.

Crespo. J. G, Goller.F, Vickers. N.J (2012). Pheromone mediated modulation of pre-flight warm up behaviour in male moths. Journal of experimental biology. 215, 2203–2209

Dangles, O., Pierre, D., Magal, C., Vannier, F., & Casas, J. (2006). Ontogeny of air-motion sensing in cricket. Journal of Experimental Biology, 209(21), 4363–4370.

Dorsett. D (1962). Preparation for flight by Hawkmoths. Journal of Experimental biology. 39, 579–588.

Dotterweich, K. (1928). Beitrflge zur Nervenphysiologie der Insekten. Zool. Jb. (AUg. Zool. PkytioL), 44, 399–425

Eaton. J. L (1971). Morphology of the head and thorax of the adult tobacco hornworm *Manduca sexta* (Lepidoptera: Sphingidae). 1. Skeleton and Muscles. Annals of the Entomological Society of America.

Esch, H. (1964). Ueber den Zusammenhang zwischen Temperatur, Aktionspotentialen undThoraxbewegungen bei der Honigbiene (Apis mellifica L.). Z. vergl. Physiol. 48, 547–55

Esch, H. (1991). Neural control of fibrillar muscles in bees during shivering and flight. J. Exp. Biol. 159, 419–431.

Field, L. H., & Matheson, T. (1998). Chordotonal Organs of Insects. Advances in Insect Physiology Volume 27, 1–228. doi:10.1016/s0065-2806(08)60013-2

Gewecke (1970). Antennae: Another wind sensitive receptor in locusts. Nature, Vol.225, 1263–1264

Gewecke, M., Heinzel, H. G., & Philippen, J. (1974). Role of antennae of the dragonfly *Orthetrum cancellatum* in flight control. Nature, 249(5457), 584–585.

Gewecke, M.(1975). The influence of the air current organs on flight behaviour of *Locusta migratoria*. J. Comp. Physiol. 103, 79–95

Guthrie. D. M (1964). Observations on the nervous system of the flight apparatus in Locust *Schistocerca gregaria*. Journal of Cell Science.: 183–201

Hedrick, T. L. (2008). Software techniques for two-and three-dimensional kinematic measurements of biological and biomimetic systems. Bioinspiration & biomimetics, 3(3), 034001.

Heinrich. B and Bartholomew G. A (1971). An analysis of pre-flight warmup in the sphinx moth, Manduca sexta

Heinrich B (1989). Beating the Heat in Obligate Insect Endotherms:The Environmental Problem and the Organismal Solutions. Amer. Zool., 29:1157–1168

Horsmann, U., Heinzel, H.G. & Wendler, G. (1983). The phasic influence of self-generated air current modulations on the locust flight motor. J. Comp. Physiol. 150, 427–438.

Ito, K., Shinomiya, K., Ito, M., Armstrong, J. D., Boyan, G., Hartenstein, V., … & Vosshall, L. B. (2014). A systematic nomenclature for the insect brain. Neuron, 81(4), 755–765.

Insausti T., and Lazzari C. R., (2000). The central projection of Cephalic mechanosensory axons in the hematophagus bug *Triatoma infestans*. Mem Inst oswaldo cruz, Rio de Janeiro, Vol. 95(3): 381–388.

Jerrold H. Zar (2011). Biostatistical Analysis. Pearson education. Inc

Kammer. A. B (1967). Muscle activity during flight in some large Lepidoptera. Journal of experimental biology. 47, 277–295.

Kammer. A. B (1968). Motor patterns during flight and warm up in Lepidoptera. Journal of experimental biology. 48, 89–109.

Kammer, A. E. (1971). The motor output during turning flight in a hawkmoth, Manduca sexta. Journal of Insect Physiology, 17(6), 1073–1086.

Krishnan A., Prabhakar S., Sudarshan S., and Sane S. P., (2012). The neural mechanism of antennal positioning in flying moths. The Journal of Experimental Biology, 215: 3096–3105.

Krishnan, A., & Sane, S. P. (2015). Antennal mechanosensors and their evolutionary antecedents. In Advances in Insect Physiology (Vol. 49, pp. 59–99). Academic Press.

Lehane M.J and Schofield, C.J (1981). Field experiments of dispersive flight by *Triatoma infestans*. Transactions of the Royal Society of Tropical medicine and hygiene, Vol 75, No.3

Linnaeus, C (1758). Systema Naturae per regna tria naturae, secundum classes, ordines, genera, species, cum characteribus, differentiis, synonymis, locis. Editio decima, reformata. Tomus I. Holmiae (Laurentii Salvii), pp 1–824.

Mamiya, A., Straw, A. D., Tómasson, E., & Dickinson, M. H. (2011). Active and passive antennal movements during visually guided steering in flying Drosophila. Journal of Neuroscience, 31(18), 6900–6914.

Mamiya, A., & Dickinson, M. H. (2015). Antennal mechanosensory neurons mediate wing motor reflexes in flying Drosophila. Journal of Neuroscience, 35(20), 7977–7991.

Miller JP, Krueger S, Heys JJ, Gedeon T (2011) Quantitative Characterization of the Filiform Mechanosensory bristle Array on the Cricket Cercus. PLoSONE 6(11): e27873. doi:10.1371/journal.pone.0027873

Orchard, I., Ramirez, J. M., & Lange, A. B. (1993). A multifunctional role for octopamine in locust flight. Annual review of entomology, 38(1), 227–249.

Pringle JWS (1957) Insect flight. Cambridge: Cambridge University Press.

Rind. F. C (1982). The organization of flight motor neurons in the moth, *Manduca sexta*. Journal of experimental biology, 102, 239–251.

Sadaf, S., Hasan, G. (2014). Serotonergic neurons of the *Drosophila* air-pulse-stimulated flight circuit. J Biosci 39, 575–583.

Sadaf, S., Reddy, O. V., Sane, S. P, Hasan, G. (2015). Neural Control of Wing Coordination in Flies, Current Biology, Volume 25, Issue 1, Pages 80–86

Sane, S. P., & Jacobson, N. P. (2006). Induced airflow in flying insects II. Measurement of induced flow. Journal of Experimental Biology, 209(1), 43–56.

Sane, S. P., Dieudonné, A., Willis, M. A., & Daniel, T. L. (2007). Antennal mechanosensors mediate flight control in moths. Science, 315(5813), 863–866.

Sane S.P, Srygley. R. B and Dudley. R (2010). Antennal regulation of migratory flight in the neotropical moth *Urania fulgens*, Biol. Lett. 6,406–409.

Sant, H. H., & Sane, S. P. (2018). The mechanosensory-motor apparatus of antennae in the Oleander hawk moth (*Daphnis nerii*, Lepidoptera). Journal of Comparative Neurology, 526(14), 2215–2230.

Savela, Markku. “*Daphnis* Hübner, [1819]”. Lepidoptera and Some Other Life Forms.

Soltavalta, O. (1954). On the thoracic temperature of insects in flight (contributions to the problem of insect flight IV.). Ann Zool Soc Zool Bot Fenn Vanamo, 16, 1–22.

Snodgrass, R. E. (2018). Principles of insect morphology. New York London: Mc Graw-Hill Book company, Inc. 514–517.

Sponberg S and Daniel T. L. (2012). Abdicating power for control: a precision timing strategy to modulate function of flight power muscles, Proc. R. Soc. B.2793958–3966

Srinivasan, M. V., Zhang, S. W., Lehrer, M., & Collett, T. S. (1996). Honeybee navigation en route to the goal: visual flight control and odometry. Journal of Experimental Biology, 199(1), 237–244.

Srygley R. B (2001), Sexual differences in tailwind drift compensation in *Phoebis sennae* butterflies (Lepidoptera: Pieridae) migrating overseas, Behavioral Ecology, Volume 12, Issue 5, Pages 607–611

Steinmann, T., Casas, J., Krijnen, G., & Dangles, O. (2006). Air-flow sensitive hairs: boundary layers in oscillatory flows around arthropod appendages. Journal of experimental biology, 209(21), 4398–4408.

Stevenson P. A, Kutch W (1987). A reconsideration of the central pattern generator concept for locust flight, J Comp Physiol A (1987) 161:115–129

Stevenson, R. D., & Josephson, R. K. (1990). Effects of operating frequency and temperature on mechanical power output from moth flight muscle. Journal of Experimental Biology, 149(1), 61–78.

Taylor B, Jones MD. (1969). The circadian rhythm of flight activity in the mosquito *Aedes aegypti* (L.). The phase-setting effects of light-on and light-off. J Exp Biol.

Trimarchi, J.R., Schneiderman, A.M. (1995). Flight initiations in *Drosophila melanogaster* are mediated by several distinct motor patterns. J Comp Physiol A 176, 355–364.

Tyrer N. M., Bacon J. P., and Davies C.A., (1979). Sensory Projections from the Wind-sensitive head bristles of the Locust, *Schistocerca gregaria*. Cell and Tissue Research, 203: 79–92.

Wolbarsht M, (2004). Electrical characteristics of insect mechanoreceptors. Journal of General physiology

Weis-Fogh (1949)-AN aerodynamic sense organ stimulating and regulating flight in locusts. Nature. 164, 873–874

Weis-Fogh, T. (1956). Biology and physics of locust flight IV. Notes on sensory mechanisms in locust flight. Philosophical Transactions of the Royal Society of London. Series B, Biological Sciences, 239(667), 553–584.

