## Supplementary methods for "Mechanosensory cephalic bristles mediate rapid flight initiation in endothermic hawkmoths"

Maitri Manjunath*[,1], Chinmayee. L. Mukunda* and Sanjay. P. Sane*
* National Centre for Biological Sciences,
Tata Institute of Fundamental Research,
GKVK Campus, Bellary Road
Bangalore 560065, Karnataka,
India

[1] SASTRA University,
Thanjavur,
Tamil Nadu,
India

**Extracellular recording from wing muscles post cephalic bristles stimulation:**

*Stimulus*

We used a precise moving coil servo motor (300C-I mechanical stimulator, Aurora Scientific, ON, Canada) with a disc attachment (see Materials and methods) to provide tactile stimulus to the cephalic bristles. Two kinds of stimulus protocols were provided.

i) One-step protocol: A single-step pulse stimulus of 0.5 seconds.

ii) Three-step protocol: A stimulus consisting of 3-step pulses of 0.5 seconds each separated by 4.5 seconds.

In both protocols, baseline activity was recorded for 2 seconds before the first stimulus followed by the 0.5 seconds response to stimulation, and a 4.5 seconds post-stimulus response. This was the repeated twice in succession for the three-step trial.

The responses of the muscles were recorded for 3 repetitions of each protocol with 25 minutes of rest between trials. This time interval was sufficient for the moths to return to their resting state after the first stimulus. All recordings which showed variations in the motor feedback signal before the onset of the stimulus were eliminated from the dataset.

All recordings were performed with a sampling rate of 10 kHz at a gain of 1000.

*Analysis*

We recorded from 8 individuals with 6 trials each from the 4 muscle groups. Of these, we used data from only those individuals which had at least 3 valid trials. Trials were considered invalid if they had unexpected variability in the baseline of stimulus feedback before the start of stimulus (which suggested a misplaced electrode), or the muscle activity before and after stimulation were indistinguishable (which suggested no response due to dislodgement of electrode). Using these criteria, we have used data from axillary muscle recordings from 5 individuals (a total of 28 trials), basalar muscle recordings from 7 individuals (a total of 25 trials), dorso-longitudinal (DLM) muscle recordings from 7 individuals (total of 29 trials), subalar muscle recordings from 6 individuals (total 28 trials) for analyses. For statistical analysis, we included only those recordings that consisted at least 3 trials per individual. This resulted in a selection of 110 trials for further analysis. The responses of the muscles were filtered using a 3rd order Butterworth bandpass filter with cut-offs at 10 and 500 Hz.

In many cases, the response following the first step stimulus in the three-step protocol extensively overlapped with the occurrence of the next step stimulus. In such cases, only the response to the first pulse of the 3-step protocol was used for analysis. Because the analysis aimed to measure latency of the response, a total length of 2.5 seconds of recording was used for all analyses which included the pre-stimulation baseline of 2 seconds and the response of the muscle till the end of the 0.5 seconds stimulation.

In most extracellular muscle recordings, both spikes and twitches were present which made the process of spike detection challenging. Although we were able to see a clear increase in activity upon stimulation of cephalic bristles, our recordings usually had a mixed population of spikes and muscle twitches of variable widths and amplitudes. This made it difficult to use a generic spike template to identify the spikes. Hence, we designed a custom algorithm to detect spikes in the muscle recordings, which is described below.

*Spike detection*

Spikes occurring in response to the stimulation were identified by template matching and false-positive spikes were algorithmically eliminated based on certain fixed criteria. Spikes were identified using a custom-written program for template matching in MATLAB based on the *findsignal* function, a built-in function in MATLAB's signal processing toolbox. Briefly, this function identifies signals matching a given template by performing a similarity search with dynamic time warping (DTW). DTW reduces the Euclidean distance between the template and the signal by stretching, compressing, and repeating samples in the template or the signal to be matched. Thus, the DTW algorithm allows identification of spikes with slightly different shapes compared to the template. Based on the position of the electrode with respect to the motor units, the spike shapes can vary. Therefore, multiple templates were required to identify all the spikes within a recording. Thus, 1-3 spike templates were manually identified for each trial.
This procedure identified many signals that resembled the specified templates. To eliminate the false-positive detections in an unbiased manner, three criteria were employed. First, we applied

thresholds (<10 µV and >5mV) to amplitude values. We used a threshold value of 30% of spike template amplitudes to filter out detections that were dissimilar to typical muscle activity and spike template amplitudes, respectively. Second, detected spikes of width <2 ms and >13 ms were discarded. Third, redundant spikes and spikes with considerable overlaps were identified based on a 5% threshold on the extent of overlap. In such cases, the detected spike with a smaller peak-to-peak amplitude was eliminated. This procedure also eliminated some of the true spikes in the muscle activity. Finally, for every spike, the peak/trough location (whichever had larger amplitude) was used for assigning spike location.

*Estimating latency*

Latency was determined to be the time delay between the onset of stimulus and the first spike. In the recordings for which no activity was observed before the stimulus, the first spike occurring 5ms after the rising edge of the stimulus pulse was considered for calculating the latency. In the recordings for which activity was observed before the stimulus, we considered spikes larger than baseline spikes for calculating latency.

*Shortcomings*

Extracellular muscle recordings often consist of muscle twitches and muscle spikes (Pringle,1949). Achieving precise differentiation between these two types of signals proves challenging. The approach employed in this study identifies signals that match with predefined templates. The Euclidean distance between the signal and the template is a measure of how closely the signal resembles a template. A larger distance between the template and the signal implies poor matching of the two and a shorter distance signifies better matching. A high tolerance (larger distances between the template and matched signal) caused more false positives and a low tolerance led to more false negatives in spike detection in our analysis. A level of tolerance was selected to minimize the number of false positives and negatives.

The complete set of codes is available at https://github.com/chinmayeelm/EMG_spike_detection.
