## Supplementary figures and images for "Mechanosensory cephalic bristles mediate rapid flight initiation in endothermic hawkmoths"

### supplementary figures

| | | Latency(ms) | | | |
|---|---|---|---|---|---|
| **Moth number** | **Trial number** | **Leg** | **Abdomen** | **Wing** | **Antenna** |
| 1 | 1 | 154 | 175 | 182 | 311 |
| | 2 | 152 | 183 | 195 | 319 |
| | 3 | 150 | 170 | 187 | 324 |
| 2 | 1 | 196 | 205 | 292 | 332 |
| | 2 | 192 | 196 | 260 | 338 |
| | 3 | 191 | 202 | 287 | 331 |
| 3 | 1 | 250 | 300 | 533 | 571 |
| | 2 | 244 | 216 | 551 | 682 |
| | 3 | 280 | 260 | 608 | 691 |
| 4 | 1 | 385 | 455 | 562 | 552 |
| | 2 | 367 | 468 | 565 | 527 |
| | 3 | 370 | 479 | 515 | 577 |
| 5 | 1 | 240 | 480 | 534 | 469 |
| | 2 | 254 | 453 | 554 | 465 |
| | 3 | 228 | 460 | 520 | 491 |
| 6 | 1 | 269 | 428 | 430 | 503 |
| | 2 | 279 | 414 | 460 | 473 |
| | 3 | 282 | 419 | 481 | 502 |
| 7 | 1 | 265 | 681 | 491 | 522 |
| | 2 | 228 | 357 | 470 | 544 |
| | 3 | 228 | 356 | 466 | 543 |
| 8 | 1 | 278 | 307 | 352 | 469 |
| | 2 | 285 | 298 | 273 | 479 |
| | 3 | 300 | 319 | 323 | 509 |
| 9 | 1 | 436 | 490 | 562 | 548 |
| | 2 | 328 | 397 | 404 | 464 |
| | 3 | 334 | 449 | 426 | 397 |
| 10 | 1 | 540 | 519 | 510 | 450 |
| | 2 | 608 | 446 | 544 | 492 |
| | 3 | 546 | 400 | 370 | 524 |

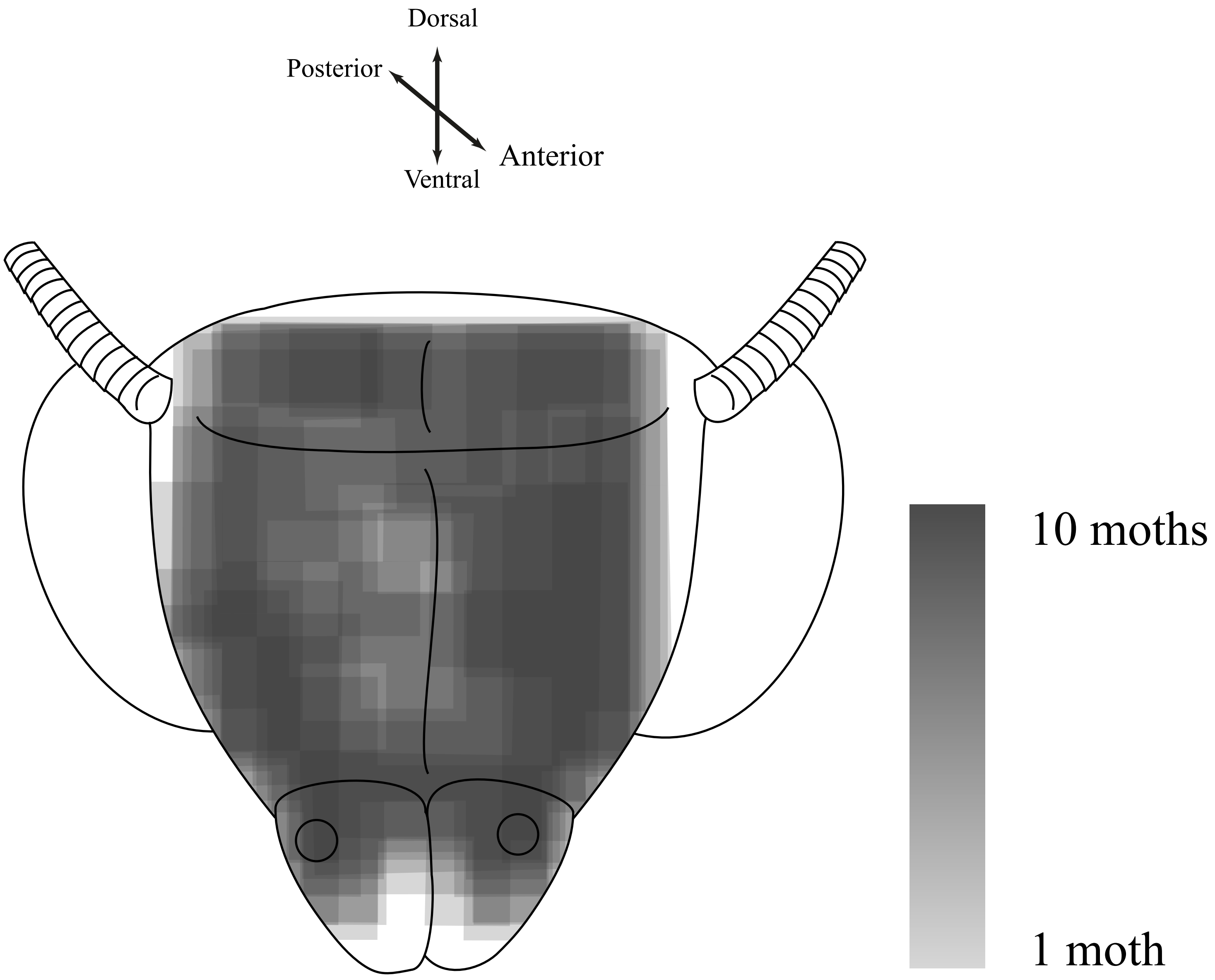


**Frontal View**

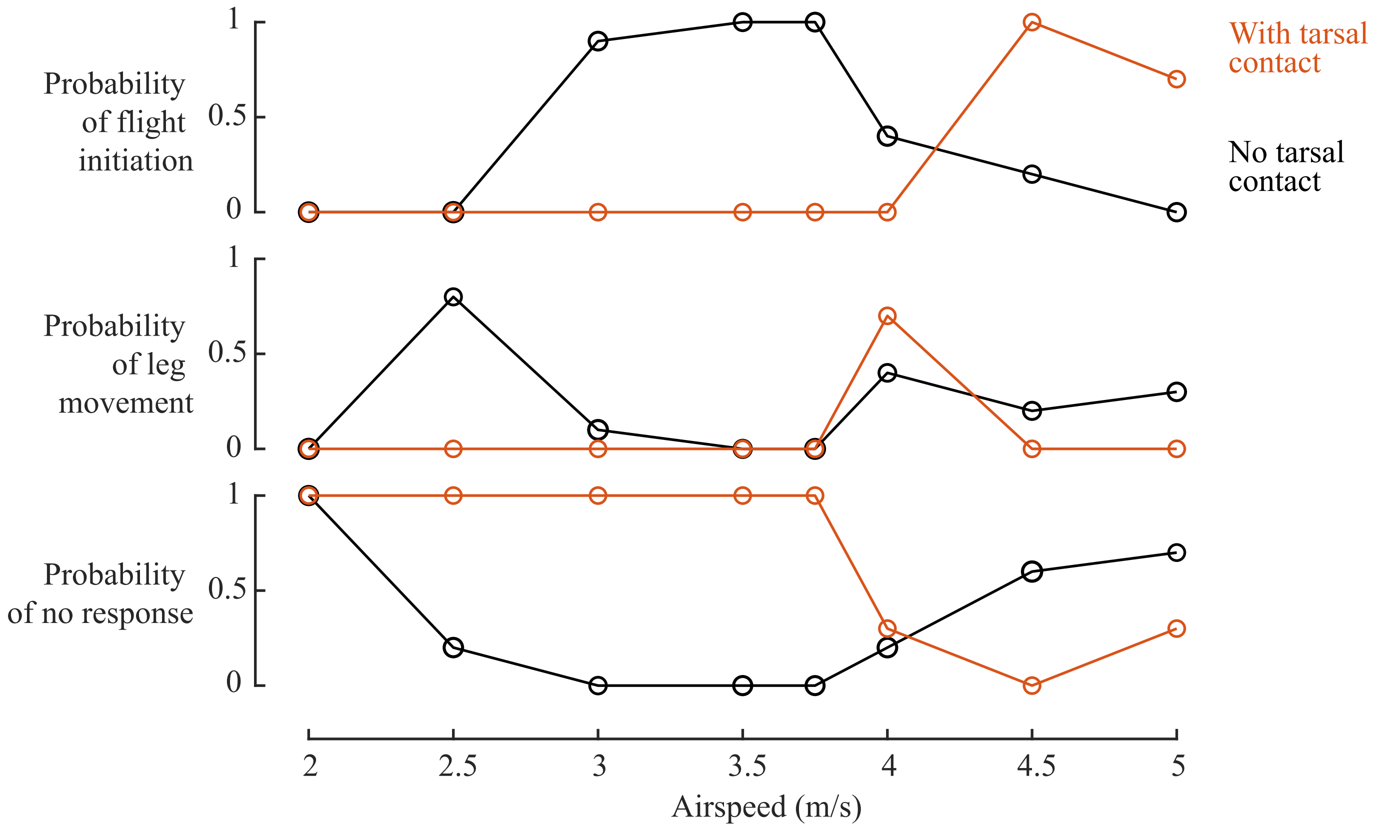
